# pSiteExplorer: Visualization of the mass spectrometric data underlying phospho-site quantification

**DOI:** 10.1101/2021.09.01.458576

**Authors:** Michael J. Plank

## Abstract

Mass spectrometry based phospho-proteomics is a widely used approach to assess protein phosphorylation. Intensities of phospho-peptide ions are obtained by integrating the MS signal over their chromatographic peaks. How individual peptide measurements mapping to the same phospho-site are combined for the quantification of the given site is, however, in most cases hidden from researchers conducting, reviewing, and reading these studies. I here describe pSiteExplorer, an R script that visualizes the peak intensities associated with phospho-sites in MaxQuant output tables. Barplots of MS intensities originating from phospho-peptides with distinct amino acid sequences due to missed cleavages, different numbers of phosphates and from all off-line chromatographic fractions and charge states are displayed. This tool will help gaining a deeper insight into phospho-site quantifications by contrasting individual and summed phospho-peptide intensities with the site-level values derived by MaxQuant. This will support the validation of quantification results, for example, for the selection of candidates for follow-up studies.

## Introduction

Phospho-proteomics has seen an increase in popularity over the last years, resulting in its wide-spread use in cell biology. In this context, researchers using, and interpreting phospho-proteomics results are frequently not the same as the ones generating the data. End-users may not always be aware of the early steps of data processing and that phospho-site quantification arises from the selection and aggregation of intensities of phospho-peptides, which constitute the actual analytes measured. Data may instead be analyzed on peptide level, alleviating most of the problems described in the following, however, the entity of biological interest usually is a phospho-site, rather than a phospho-peptide.

Therefore, the analysis of phospho-proteomic data in the context of the biological question under study generally starts from a table listing one quantitative value per phospho-site per sample. At this point, it is no longer obvious that frequently several integrated peaks, potentially corresponding to peptides of differing sequence, underly a single phospho-site quantification. It is also hidden from the end-user that signals originating from a phospho-site of interest may be ignored for the quantification of the site by the analysis software based on a number of criteria.

Challenges inherent to bottom-up phospho-proteomics workflows, such as the existence of multi-phosphorylated peptides, uncertainties in phospho-site localization, and co-elution of positional isoforms may in turn lead to problems in translating measurements on the phospho-peptide level to quantification on the level of phospho-sites.^1^ To illustrate a problem that may arise, consider two phospho-sites P1 and P2, located in close proximity to each other, of which P2 is phosphorylated upon a stimulus. As P1 and P2 reside on the same phospho-peptide, the peptide singly-phosphorylated on P1 decreases upon stimulation due to phosphorylation of P2. If therefore the doubly phosphorylated peptide is not taken into account for the quantification of P1, researchers may erroneously report that the protein is dephosphorylated at P1 upon stimulation.^2^

I here introduce pSiteExplorer, an R-script for the visualization of data underlying phospho-site quantification. pSiteExplorer presents the signals contributing to the quantification of a site of interest, to reveal the measurements that have been excluded in the process and to provide a better understanding of how peptide-level data are filtered and combined during the analysis process.

pSiteExplorer is written in the R-language and utilizes output from the MaxQuant software as input. MaxQuant is a widely used data analysis tool used in proteomics due to its power and free availability. Important to this study, MaxQuant provides outputs both on the phospho-peptide and phospho-site level. In its current version, pSiteExplorer utilizes data from data-dependent mass spectrometry experiments employing precursor-labelling (e.g. SILAC) and supports data generated using offline fractionation.

## Methods

### Implementation and overview

pSiteExplorer is a program written in R-code ^3^ to provide visualization of quantitative MS data underlying the quantification of selected phospho-sites on selected proteins. The program code is provided as a supplement to this manuscript. Usage requires installation of the required code packages reshape2 ^4^, ggplot2 ^5^, stringr and Biostrings ^6^. Execution of the program can easily be accomplished with minimal user experience.

The program generates barplots depicting label ratios and normalized intensities (in the following simply referred to ‘intensities’) on phospho-site level and of phospho-peptide intensities associated with the corresponding site. The application of pSiteExplorer is currently limited to mass spectrometry data generated by data dependent acquisition using precursor-labelling (e.g. SILAC ^7^). The program is intended for manual inspection and validation and handles one phospho-site at a time.

### User input

User input to pSiteExplorer is provided by supplying the necessary information in lines 33 - 46 of the R-script. A readMe-file associated with the program provides detailed instructions for the required input.

pSiteExplorer produces visualizations from output of the MaxQuant software. MaxQuant is a freely available software tools and one of the most widely used programs for the analysis of data-dependent mass spectrometry experiments. ^8^ Of the output tables written by MaxQuant, which allow interrogation of the results from different perspectives and on different levels, pSiteExplorer requires the ‘Phospho (STY)sites.txt’ and ‘Evidence.txt’ tables. ‘Phospho (STY)sites.txt’ is generated if ‘Phospho (STY)’ is specified as a variable modification in the database search and lists one phospho-site per row. Intensities and label-ratios constitute aggregates from all corresponding phospho-peptides passing the applied filtering criteria from all fractions and charge states. The ‘evidence.txt’ file lists evidences of identified peptides. An ‘evidence’ is a MS/MS identification mapped to a feature multiplet (i.e. corresponding features from all SILAC-channels) or, in some cases, an MS/MS-ID without associated MS1 signal. MS/MS identifications can also be associated with features via match-between-runs.

In addition to the file location of the stated MaxQuant output, the phospho-site and name of the protein of interest as it appears in the ‘Protein’ column of ‘Phospho (STY)sites.txt’, the sample names as given in the MaxQuant ‘Experiment’ input column are required. Further, the user is asked to provide freely chosen names for the experimental conditions studied and an experimental design of how these conditions are associated with SILAC-channels in each sample. The user is also required to supply the name of the fasta file used for the database search, which needs to be located in the specified file location.

Optionally, it can also be defined if y-axes of plots should be scaled to the highest value locally or globally (local scaling is used by default). Further, a minimum site-localization score can be defined. This value sets a threshold above which data should be considered when displaying signals not qualifying as evidences in the MaxQuant algorithm (see below) and to serve as a threshold in plot annotations. The default value is 25%. This parameter does not affect the selection of signals as evidences, as this is inherent to MaxQuant. Finally, there is an option to alter the resolution of plots.

### Algorithm

pSiteExplorers obtains intensities for the phospho-site of interest from the ‘Phospho (STY)Sites.txt’ MaxQuant output file, arranges them according to the user-defined experimental design and produces a barplot.

The evidences contributing to the quantification of the site are extracted from the evidence.txt file by looking up its ‘Evidence IDs’. pSiteExplorer collects the evidences for the phospho-site of interest and divides them into isoform-groups. An ‘isoform-group’ is defined as a group of peptides with the same base-sequence and number and type of modifications. To be listed as an evidence for a phospho-site by MaxQuant, the evidence needs to exceed MaxQuant`s localization score threshold. The grouping into isoform-groups permits aggregation of positional isomers with respect to other modifications than the phospho-site under investigation. An alternative approach would be to aim for the quantification of corresponding features linked across samples. The design choice for the use of isoform-groups was made to avoid imposing an arbitrary localization score threshold and separating features from different samples that fall to different sides of the threshold due to randomness. Also, as positional isoforms of multiple-modified peptides often overlap chromatographically, a clean linking of corresponding features is not always possible.

Intensities from different evidences mapping to an isoform-group are added up within each fraction, charge-state, sample and labelling-channel. A barplot is produced per fraction and charge state in which phospho-peptides of the isoform-group were observed.

As stated above, identifications need to pass MaxQuant`s localization score threshold to be considered evidences for a phospho-site. To allow the inspection of additional ‘unused’ features, with higher localization ambiguity, pSiteExplorer searches for additional features above a user-defined localization score threshold. The features are obtained by searching ‘evidence.txt’ for sequences from the protein of interest that overlap the phospho-site of interest based on the user-supplied fasta-file.

### Output

Even though MaxQuant also provides label-ratios for SILAC data, pSiteExplorer displays intensities on evidence-level, as they provide a layer of information that is hidden when looking at ratios, most notably, if a peptide contributes strong signal or is closer to background level.

pSiteExplorer displays phospho-site intensities as reported in ‘Phospho (STY)sites.txt’ by MaxQuant (“pSite – MQ”) and three additional types of plots on the phospho-site level and the level of evidences, with the latter divided into signals from isoform groups. The types of plots are shown in Fig. 1.:

**Figure 1.**
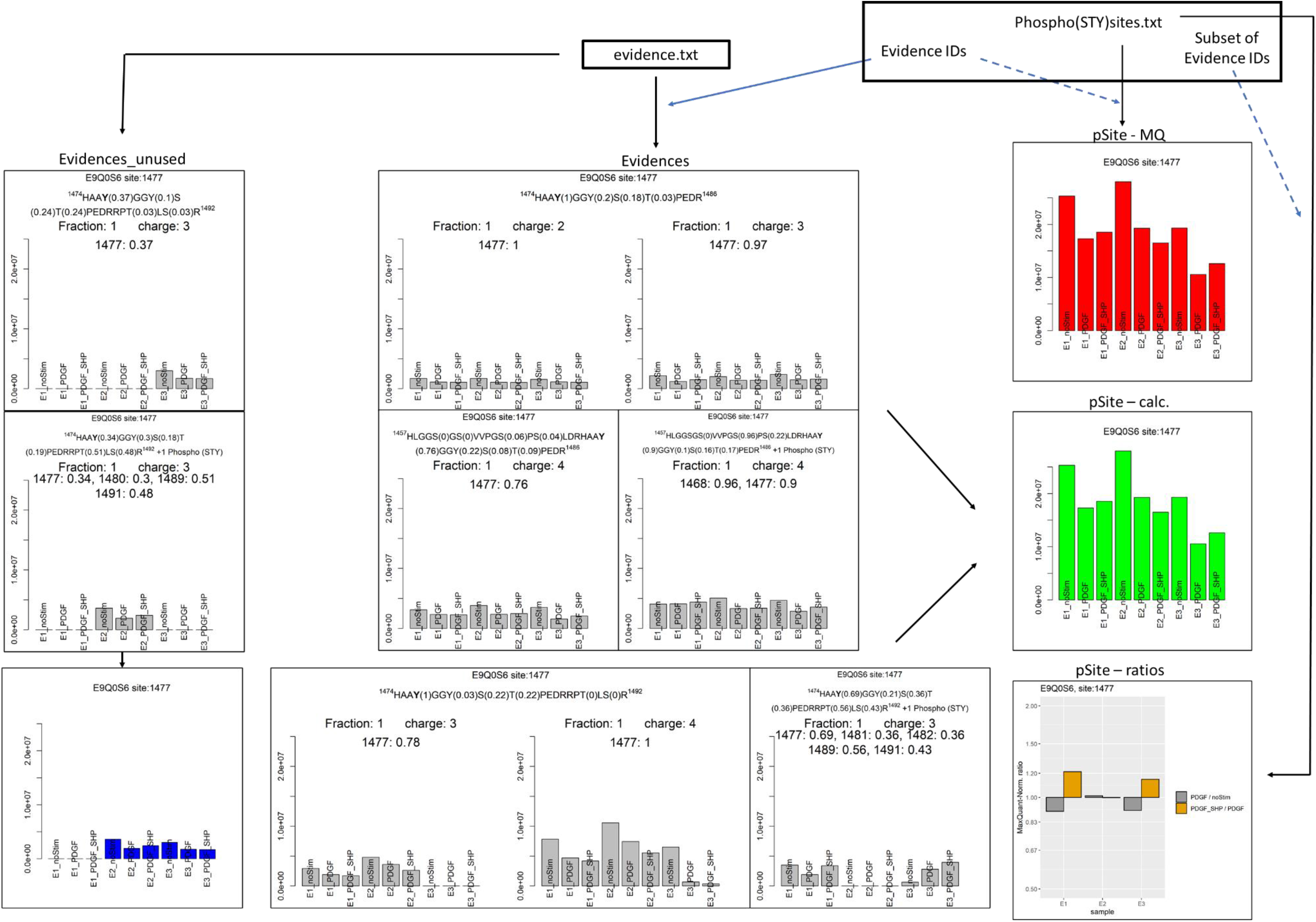
pSiteExplorer output: Plots of type ‘Evidences’ (5 isoform groups) and ‘Evidences_unused’ (2 isoform groups) are generated from evidence.txt. The former is based on signals selected as evidences for the given phospho-site by MaxQuant, while the latter displays signals not used as evidences. The ‘pSite – MQ’ plot depicts intensities as given in ‘Phospho (STY)sites.txt’. Intensities in ‘pSite – calc’ are the sums of intensities in ‘Evidences’ plots. ‘pSite – ratios’ plots depict label ratios from ‘Phospho (STY)sites.txt’. Blue lines indicate selection of data based on Evidence IDs. Full lines are selections made within pSiteExplorer and dashed lines within MaxQuant processing.

“Evidences”: A grid of barplots of summed intensities of evidences for each isoform-group. The grids are sub-divided into offline fractions and charge states.

“pSite – calc”: A barplot representing the intensities for the selected phospho-site by summing over all isoform groups in A.

“pSite – ratios”: A plot depicting the intensity ratios between channels for each sample for the given site as reported in ‘Phospho (STY)sites.txt’ by MaxQuant.

In addition, a plot of type “Evidences_unused” is produced to allow the user to explore additional evidences that include the phospho-site of interest, but are not employed for phospho-site quantification by MaxQuant. The output filename ends with the suffix “_unused”. In the cases tested, the most prevalent reason for not considering signals was that the phosphate localization probability was too low. A user defined localization score threshold (default: 25%) can be provided as a filter for evidences included in this plot.

A dummy-plot without bars is produced for isoform groups that are identified on MS/MS-level, but do not have MS-intensities associated with them.

“pSite – calc” plots are created with green bars and the output file is labelled with the suffix “_site calc”. The site-intensities reported by MaxQuant are displayed as plots with red bars and labelled with the file-suffix “_site”. This plot is intended as a sanity-check that feature intensities add up as expected, and the bar heights should be identical to “pSite – calc”. A barplot of sums of “unused” intensities is produced in addition and depicted with blue columns.

Each plot states the selected protein and site of interest in its header. Plots on evidence-level, also state the peptide sequence in the given isoform group in the grid-header. The phospho-site of interest is in bold. Each position with a non-zero localization probability is followed by this probability. The highest of these probabilities in all peptides used for the plot is given. The sum of probabilities may therefore exceed 1 (or 2 in case of a doubly phosphorylated peptide, etc.). Finally, additional modifications on the peptide, in addition to the phosphorylation of interest, are stated.

The headers of each individual plot in the grid give the minimal localization probability of the site of interest over all evidence features used for the plot, as well as chromatographic fraction and charge state. Note therefore that localization probabilities are explored from two different angles: Subplots indicate the minimal observed localization probabilities to provide an indication of the lowest-quality identifications contributing to the plot, while grid headers provide the maximum probabilities to indicate how likely a peptide with (additional or incorrect) localization to a given site may have contributed to the quantification.

## Results

### Dataset

In the following, I describe some examples of phospho-site exploration by pSiteExplorer. The underlying data are obtained from a triplex-SILAC phospho-proteomics study employing off-line fractionation ^9^. Samples for this study were prepared from mouse cells that had been treated with platelet derived growth factor (PDGF) or a combination of PDGF and Shp2-inhibitor or unstimulated control cells. MaxQuant output files for this study were obtained from the ProtomeXChange repository at PXD005803 ^10^.

### Example 1

One of the most apparently differentially phosphorylated sites in this study is Y29 on Rasa1 (Uniprot ID: E9PYG6). In the phospho-site quantification, the three recorded replicates agree well, in that the sample treated with PDGF and SHP-inhibitor exhibits the highest level of phosphorylation, followed by the PDGF-treated and finally the untreated sample, both with respect to reported intensities and ratios (fig. 2 a). Exploration of the ‘evidence.txt’ MaxQuant output shows that evidences for the site are present in the form of a long, N-terminally acetylated phospho-peptide, which is present in methionine-oxidized and non-oxidized form (fig. 2 a). Both are found in charge states 3 and 4 in chromatographic fraction 1. All of these evidence features agree well with the phospho-site quantification with respect to their relative intensities and therefore lend high confidence to the phospho-site quantification.

**Figure 2.**
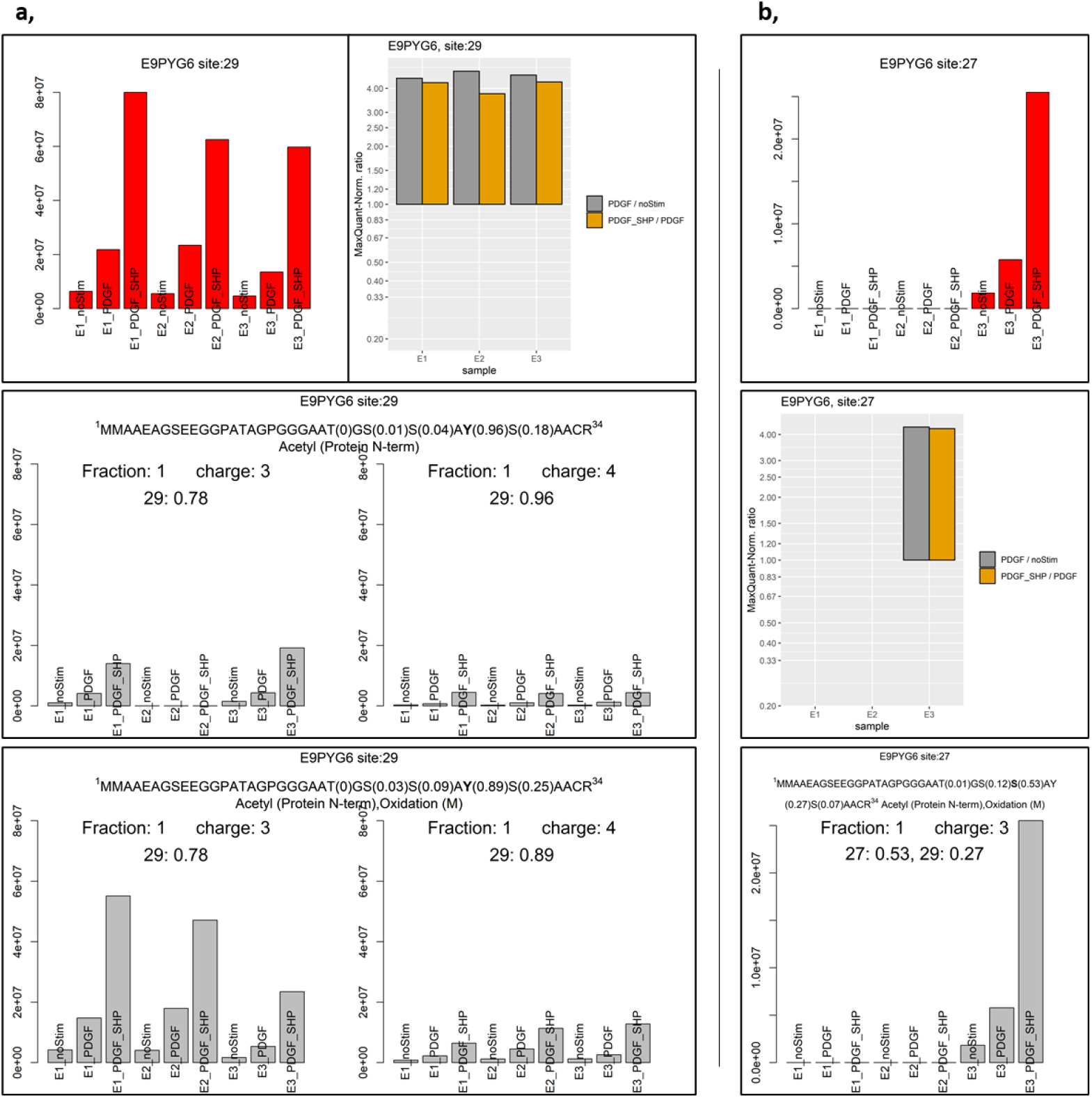
pSiteExplorer output for quantification of Y29 and S27 of protein E9PYG6. a, Quantification of Y29. Top left: Total phospho-site intensities; top right: label ratios; middle and bottom: isoform groups in two charge states each. b, Quantification of S27. Top: Total phospho-site intensities; middle: label ratios; bottom: isoform group in single observed charge state. Plots for intensities of features not used as evidences for phospho-sites are omitted.

For the close-by phospho-site S27, intensities were only recorded for the third replicate (fig. 2 b). It is found that these intensities stem from only the oxidized phospho-peptide with charge 3. The relative intensities for this replicate closely reflect the ones for Y29. The localization score for S27 is higher than the one for Y29 and slightly above 50% for this feature. Nonetheless, in a situation in which this site were of high importance for conclusions drawn from the study, this localization should be confirmed and potential co-elution with the peptide phosphorylated on Y29 further investigated.

### Example 2

An important situation in which the inspection of phospho-site quantification is advised, is when a peptide is detected with different numbers of phosphate-groups. As a second example, fig. 3 depicts signals from isoforms of singly (fig. 3a) and doubly (fig. 3b) phosphorylated peptides containing phosphorylated S213 of protein E9Q616. Peptides with additional phosphorylation of S211, S217 and S219 all contribute to the signal from doubly phosphorylated peptides. A version of the plots in which the option to plot all intensities on an equal scale was selected is shown in fig. S1.

**Figure 3.**
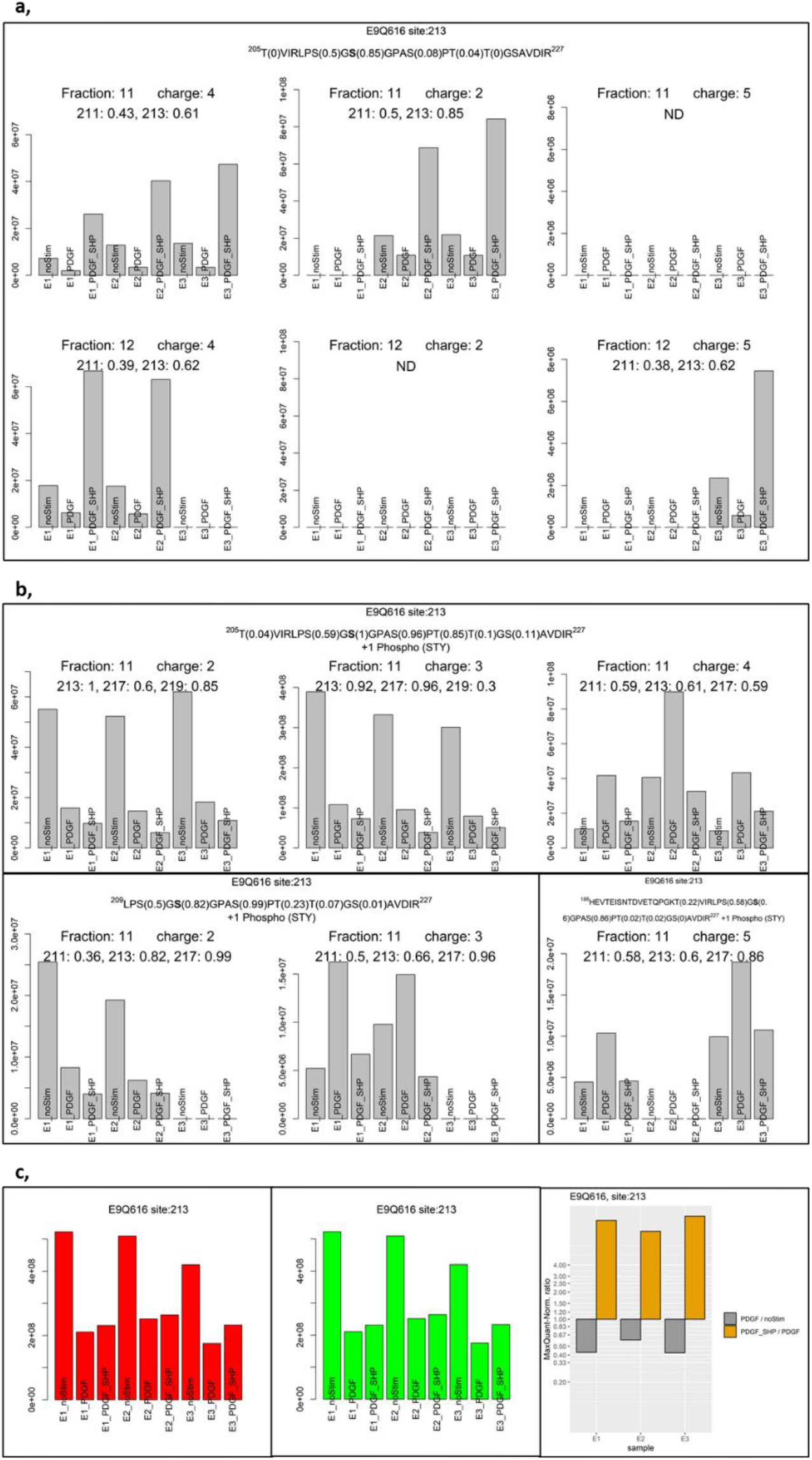
pSiteExplorer output for E9Q616 phoshorylated on residue 213. Intensities for a, singly and b, doubly phosphorylated peptides. Additional peptides which were only detected in a single replicate are omitted. c, Intensities reported by MaxQuant and calculated by pSiteExplorer and label ratios.

The singly phosphorylated isoform TVRLPSG**S**(ph)GPASPTTGSAVDIR is present in charge states 4 and 2 in fraction 11 and with charge 4 and 5 in fraction 12. A consistent moderate decrease in signal from unstimulated (‘noStim’) to PDGF-treated (‘PDGF’) sample and a strong increase in the doubly-treated (‘PDGF_SHP’) sample is consistently observed. In some cases, the light-medium-heavy-triplet was only detected for a subset of the tree replicates.

In contrast, doubly phosphorylated peptides of the same base sequences (defined as amino acid sequences without considering modifications) containing S213 exhibit two distinct patterns. Pattern A, where the abundance is strongly decreased in the PDGF and further slightly in the PDGF_SHP sample. And a second (pattern B) in which the abundance is highest in the PDGF and similar in the other two samples. Notably, the pattern of the doubly phosphorylated peptide differs not only between isoform groups, but even for different charge states within the given group.

Due to the presence of high intensity versions for both patterns and the high reproducibility between replicates, it is likely that features with the given relative intensities did not simply arise from random variation. As isoform groups aggregate isoforms with potentially different phospho-site localizations, it is likely that alternative isoforms dominate the two distinct patterns.

Peptides phosphorylated only on S211 reveal a pattern that resembles that of peptides phosphorylated only on S213, while evidences of doubly phosphorylated peptides for S211 resemble pattern B of S213 (Fig. S2).

Evidences for doubly phosphorylated peptides containing S217 follow the same pattern B, while its singly phosphorylated pattern is distinct (Fig. S3). Interestingly, the pattern for evidences of doubly phosphorylated peptides containing T219 resemble pattern A of S213 (Fig. S4). Peptides with a single phosphate localizing to T219 with a sufficiently high localization score were not detected.

The site intensities reported in ‘Phospho (STY)sites.txt’ for S213 agree, as expected, with the summed intensities of the corresponding evidences, as shown in Fig. 3C. (This is more apparent when displaying all plots on the same scale as in Fig. S1C). The higher phospho-site intensity of the ‘noStim’ sample is mainly explained by the doubly phosphorylated peptide. A different picture emerges when looking at the label-ratios reported by MaxQuant (Fig. 3C), where the intensity of the ‘PDGF’ sample is about 2x lower than the ‘noStim’ sample and of the ‘PDGF_SHP’ sample more than four times higher than of the ‘PDGF’ sample. These patterns are explained by the fact that MaxQuant takes into account all evidences for reporting intensities, but, if available, only evidences from singly phosphorylated peptides for label-ratios.

While, taken for themselves, the intensities and ratios reported for S213 on E9Q616 on phospho-site level therefore suggest a clear and consistent response, the actual level of phosphorylation at the given site may differ dramatically, depending on the prevalence of the doubly-phosphorylated peptide. Consideration of data from all versions of a phospho-peptide is therefore advised in the interpretation of changes observed for a phospho-site.

## Discussion

I here introduced pSiteExplorer, a program to visualize quantitative mass spectrometry data underlying phospho-site quantification.

A first application of pSiteExplorer is the evaluation of how well the quantifications of phospho-peptides harboring the same phospho-site, but with e.g. differing numbers of missed cleavages or from different charge states, agree with each other. In visualizing all phospho-peptides associated with a phospho-site, the program allows for a straightforward overview of the base sequence(s) the phospho-site of interest was observed on and if their quantitative patterns reflect each other. This step may prove informative as to whether, for example, the phosphorylation impacted proteolytic cleavage. Further, it can be easily evaluated if quantitative profiles agree well over replicates, charge states and off-line fractions. Finally, pSiteExplorer provides separate visualizations for phospho-peptides harboring the site of interest, but not considered as ‘evidences’ for the site by MaxQuant, usually due to ambiguous site localization. Evaluation of these ‘unused’ data may prove useful if their abundances and/or reproducibility between replicates exceeds that of the phospho-peptides used for site-quantification. Attention to these data may also be warranted if the main expected sequence, generally the one without missed cleavage and additional modifications, is only present among the unused data. In this scenario, an alternative acquisition strategy that allows for better site localization on the peptide of interest may be necessary.

Further, pSiteExplorer output depicts with which additional modifications (phosphorylation and other modifications specified in the database search) the phospho-site of interest was detected. As exemplified above, this may give clues if differential abundance of modifications at neighboring sites may be (partially) responsible for the quantification at the phospho-site level. The scope of pSiteExplorer is to provide an as complete as possible picture of available data on phospho-peptide level, but it does not aim to provide an interpretation. In cases in which differences in abundance observed on a singly phosphorylated peptide are reflected by those of multi-phosphorylated versions, it can be confidently assumed that the differences are indeed due to differential phosphorylation at the site of interest. If this is not the case, inspection of phospho-peptide data of neighboring sites is advised, but an unambiguous conclusion may be precluded by incompleteness of the data, co-elution of positional isomers and ambiguities in site localization. Targeted acquisition of all relevant species with quantification of site-determining ions on MS/MS-level may be warranted in this scenario. While the exact position may not always be critical for the molecular consequence of protein phosphorylation, its localization may be helpful in deciphering the upstream regulation, e.g. via a kinase motif. A second main application of pSiteExplorer is to evaluate how well phospho-peptide quantifications agree with the reported phospho-site quantification. In this context, the program may also prove useful in obtaining a better understanding of the algorithms applied by MaxQuant. pSiteExplorer displays data on the phospho-site level both as intensities and label-ratios. Exploring the examples provided in this manuscript and others, it became apparent that different criteria are used for the inclusion of phospho-peptides in these two measures. These somewhat depend on the processing parameters selected in MaxQuant. In the example explored here, MaxQuant apparently summed all evidences, including those of multiple-phosphorylated peptides, into phospho-site intensities, while only those of singly phosphorylated peptides contributed to label-ratios. The two measures may therefore vastly disagree, depending on the detection of multi-phosphorylated peptides and their disagreement with singly-phosphorylated ones. It may be informative to explore which phospho-peptides are and which are not in agreement with the overall phospho-site quantification. Quantitative phospho-proteomics studies employing MaxQuant for the processing of SILAC experiments naturally report label ratios, rather than intensities, on phospho-site level. Ignoring multiple-phosphorylated peptides in this measure, overall appears to be a sensible choice from a standpoint of attributing phosphorylation changes to the correct site, as abundance changes in a muliple-phosphorylated peptide are equally likely to arise from differential phosphorylation at the other sites. This strategy may however miss for example a situation in which a site is only phosphorylated upon a stimulus if a neighboring site is already phosphorylated, as is the case in the requirement for priming phosphorylation for GSK-3 ^11^. Also, as exemplified in the introduction, altered abundance of singly-phosphorylated peptides can potentially arise from phosphorylation changes at neighboring sites, so that ignoring multiple-phosphorylated peptides may lead to erroneous conclusions in this case ^2^. Exploring phosphorylation patterns on the phospho-peptide level using pSiteExplorer can prove useful to detect such special circumstances.

## Supporting information

readMe

pSiteExplorer R script

## ASSOCIATED CONTENT

The following files are available free of charge:

pSiteExplorer R-script; read-me file with instructions for use; data of Fig. 3 displayed with equal y-axis scales (Fig. S1); Visualization of evidences for phospho-site 211 on E9Q616 (Fig. S2); Visualization of evidences for phospho-site 217 on E9Q616 (Fig. S3); Visualization of evidences for phospho-site 219 on E9Q616 (Fig. S4).

## ACKNOWLEDGEMENT

I thank Prof. Arminja Kettenbach (Dartmouth College) for critical reading of the manuscript.

## ABBREVIATIONS

SILAC: Stable isotope labelling with amino acids in cell culture
PDGF: platelet derived growth factor

## Supplementary Figures

**Figure S1.**
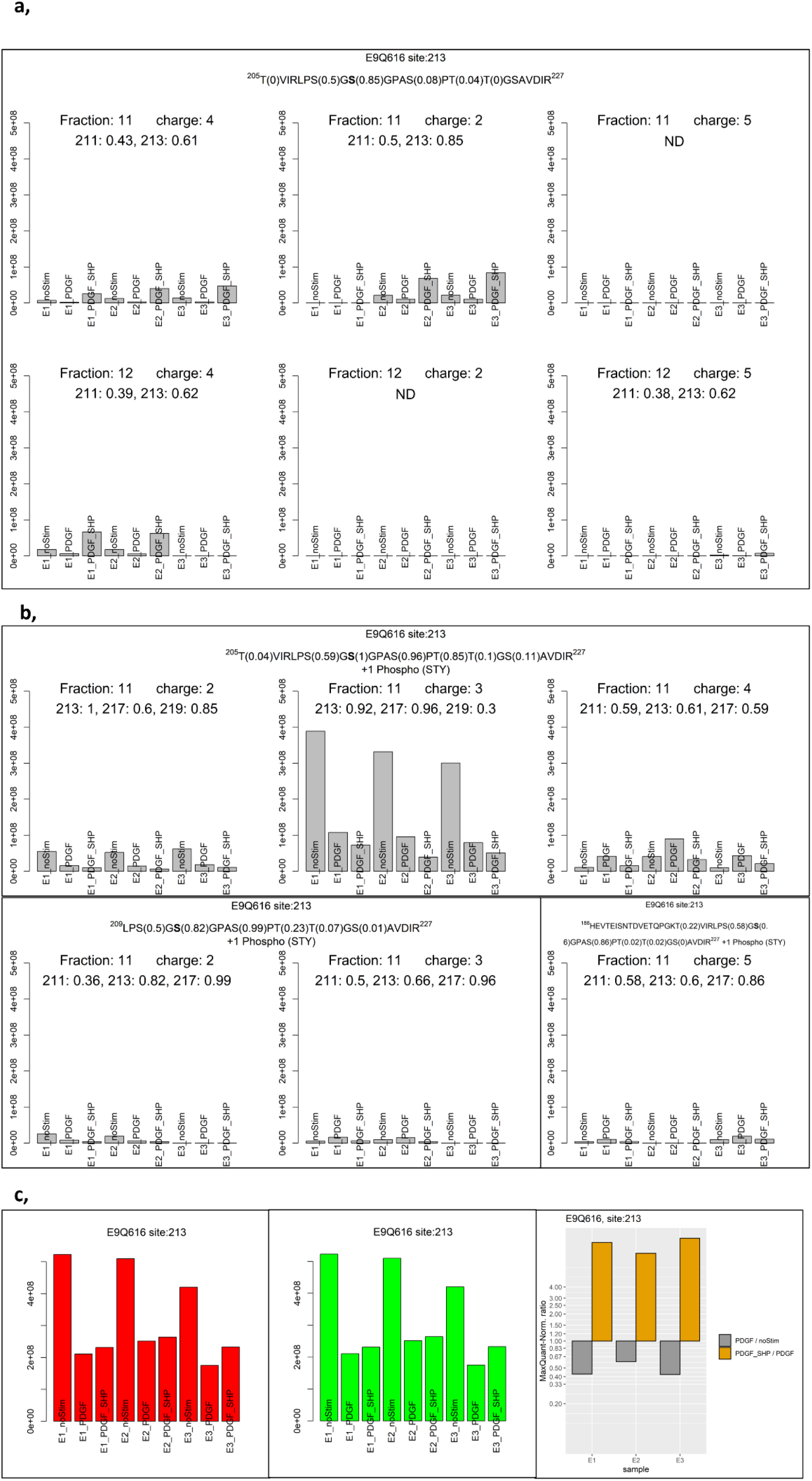
Visualization of data in fig. 3 with equal Y-axis scales. pSiteExplorer output for E9Q616 phoshorylated on residue 213. Intensities for a, singly and b, doubly phosphorylated peptides. Additional peptides which were only detected in a single replicate are omitted. c, Intensities reported by MaxQuant and calculated by pSiteExplorer and label ratios.

**Figure S2.**
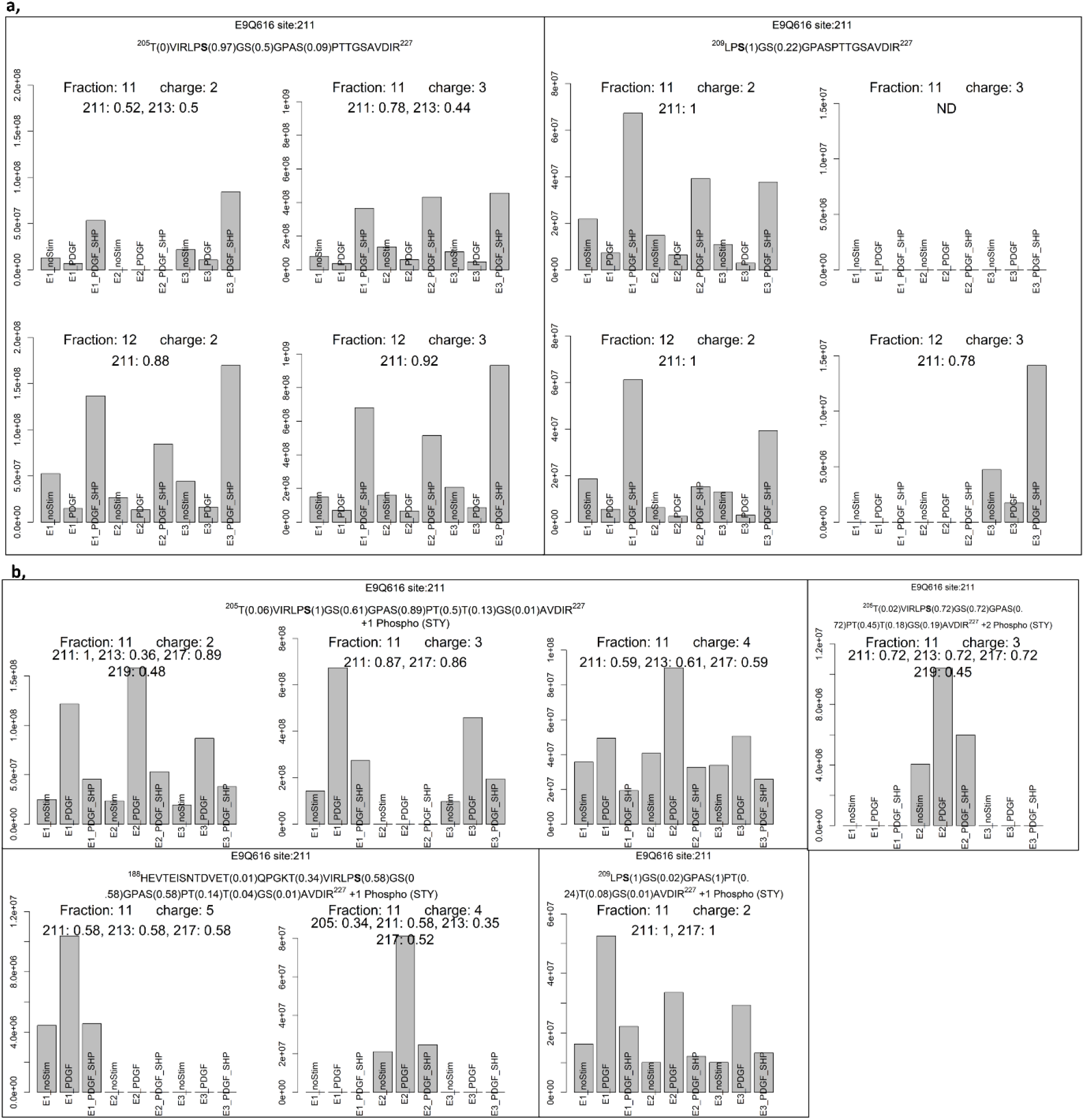
Visualization of evidences for phospho-site 211 on E9Q616. Signals from a, singly and b, doubly phosphorylated peptides are shown.

**Figure S3.**
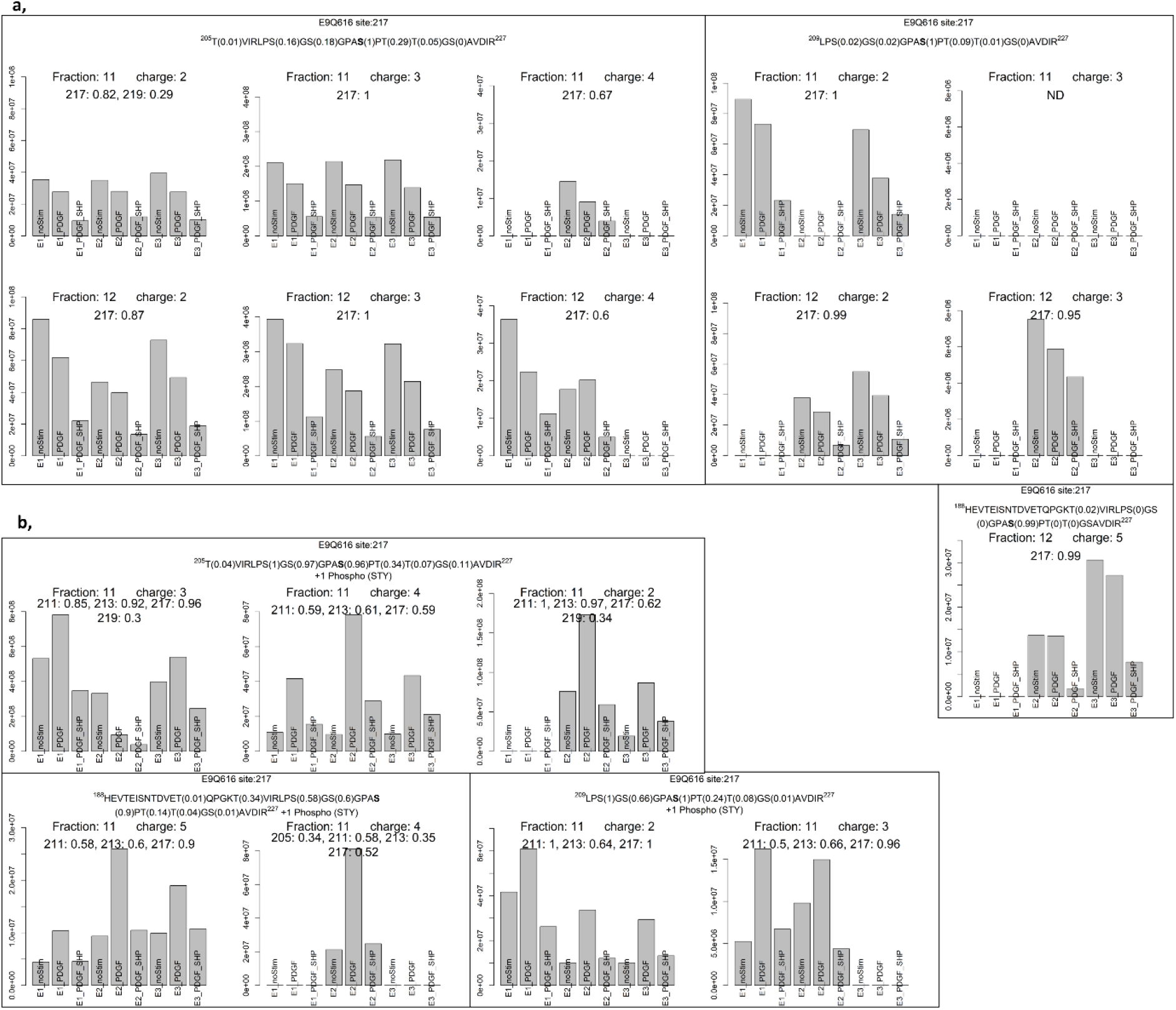
Visualization of evidences for phospho-site 217 on E9Q616. Signals from a, singly and b, doubly phosphorylated peptides are shown.

**Figure S4.**
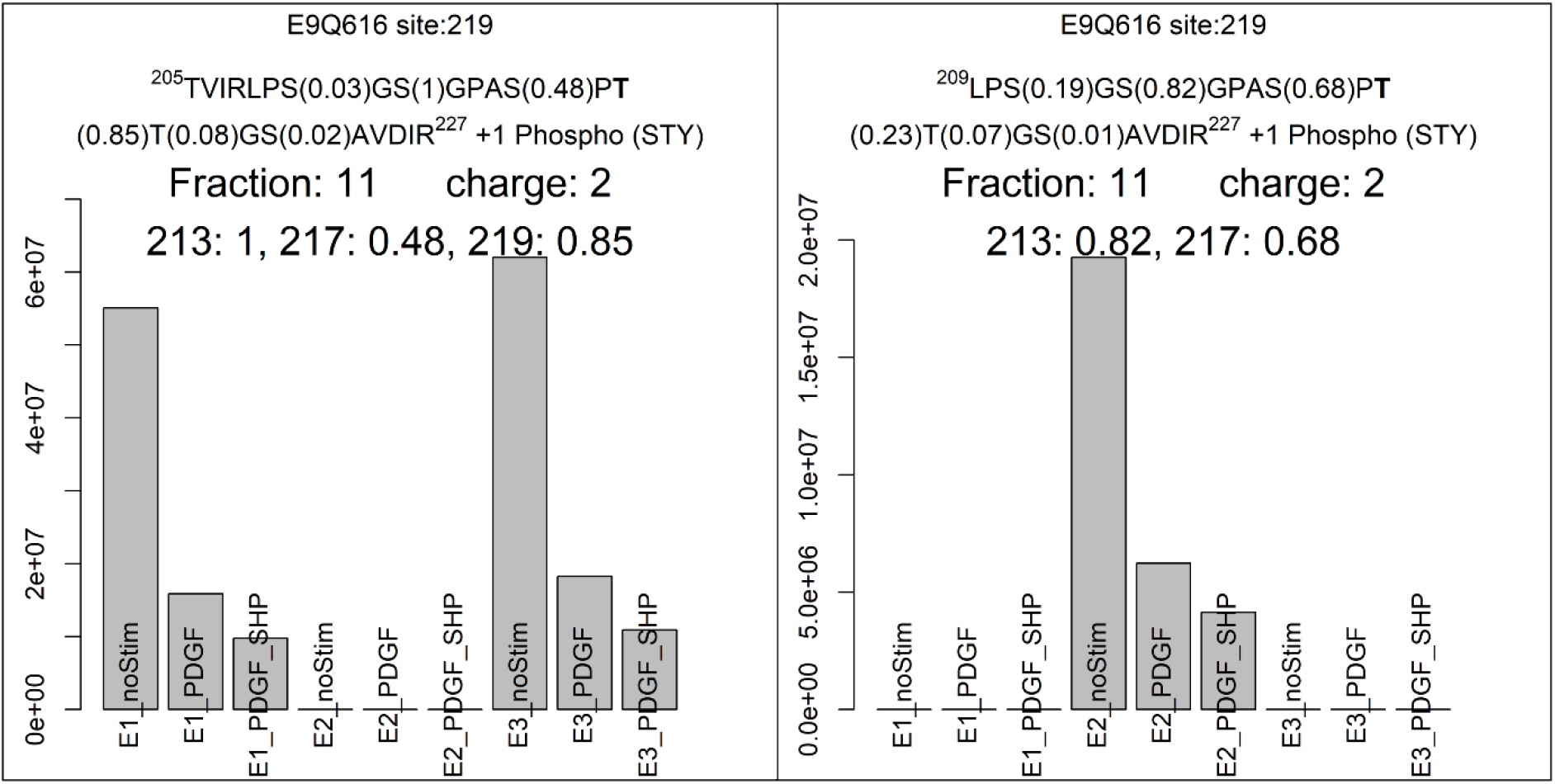
Visualization of evidences for phospho-site 219 on E9Q616.

