## Supplementary material for "pSiteExplorer: Visualization of the mass spectrometric data underlying phospho-site quantification": readMe

pSiteExplorer.R is an R script that facilitates the visualization of mass spectrometric signals associated with phospho-sites of interest.
 The program in its current version only supports precursor-labelled/SILAC data.

The program requires packages 'reshape2', 'ggplot2' and 'stringr' as well as Bioconductor 'Biostrings' package.

As input, provide 
  MaxQuant output files 'Phospho (STY)Sites.txt' and 'evidence.txt' and
  the fasta-file used for the MaxQuant search
in the same folder.

Provide the following information on user input (lines 33 to 46):
 -'file_location'   pointing to the location of input files; note that R uses '/' on Windows
 -'protein_oI'      protein of interest as in 'Protein' column in 'Phospho (STY)sites.txt'
 -'site_oI'         phospho-site of interest as 'Position within proteins' in Phospho (STY)sites.txt
 -'samp_names'      sample names as in MaxQuant "Experiment" definition and as used in column names of MaxQuant output files
 -'conds'           experimental conditions; user chooses names
 -'expDes2'         experimental design: a matrix with SILAC-labels in order of conds (columns) in each sample (rows); e.g. for a one sample with conds 
                    #in order L-M-H and second sample in M-H-L, use 'c(c('L','M','H'),c('M','H','L'))'
 -'same_scale'		determines if all plots should be displayed on the same y-axis scale (y) or normalized to the maximum within the plot (n)
 -'fasta'			filename of the .fasta file used for MaxQuant search
 -'minLoc'			minimum localization probability for displaying evidences not used for site-quantification by MaxQuant; also indicates lower
                    #threshold for localization probabilitites to annotate in figure headers
					
Expected output:
 .png files of the form '20200214_HS90B_226_isofGr_1.png', where the filename consists of date(yyyymmdd)_protein name_site_isoform group number;
  an isoform group contains phospho-isoforms, i.e. same sequence and number of phosphates with potentially alternative localization.
  The file contains a barplot for each fraction and charge state in which signal for the phospho-peptide was detected;
  In the header, the phospho-site of interest is bold and associated with the maximum observed localization probability in this isoform group.
  
Found a bug? Please report it to
