## Supplementary material for "pSiteExplorer: Visualization of the mass spectrometric data underlying phospho-site quantification": pSiteExplorer R script: pSiteExplorer.R

```
rm(list=ls())
cat("\014")  #is the code to send ctrl+L to the console and therefore will clear the screen
dev.off()
plot.new()

library(reshape2)
library(ggplot2)
library(stringr)
if (!requireNamespace("BiocManager", quietly = TRUE)) {
  install.packages("BiocManager")
  BiocManager::install(version = "3.10")
}
#if not already installed:
#BiocManager::install("Biostrings")
library("Biostrings")

today= Sys.Date()
today=gsub("-","",today)

##########################################################################
##########################################################################

#Program to visually explore the data ('evidences') associated with a selected phosphosite
#'Phospho (STY)sites' samples as columns, fractions combined - only on site level
#'modificationSpecificPeptides' samples as columns, fractions combined - isoforms combined; no info about localization on peptide
#'peptides' samples as columns, fractions combined - different modified forms of peptide combined
#'allPeptides' samples as rows, fractions not combined
#'evidence' samples as rows, fractions not combined

### The program currently only handles SILAC data!

###################   USER DEFINED: #############
file_location= "C:/Users/wrtlb/Dropbox/201912_AfterPostDoc/22_PhosphoSiteExplorer/Batth2018/MaxQuant-PDGF-SHP099-SILAC"  #define site you want to inspect
protein_oI= 'E9Q0S6'   #protein of interest as in 'Protein' column in 
#Phospho (STY)sites.txt
site_oI=  1477     #phospho-site in protein as 'Position within proteins' in Phospho (STY)sites.txt
samp_names= c('E1','E2','E3')  #as in MaxQuant "Experiment" and output file column names
conds=c('noStim','PDGF','PDGF_SHP')  #conditions: user chooses names
expDes2= matrix(c(c('L','M','H'),c('L','M','H'),c('L','M','H')),byrow=TRUE,
                ncol=3)  #experimental design: labels in order of conds (columns) in each sample (rows)  
same_scale= 'y'  #Do you want all plots on same scale? ('y'/'n')
fasta= "20210410_Mmusc_uniprot_UP000000589.fasta" #make sure this is in file_location

minLoc = 25  # % ;minimum localization score on site-of-interest to include ppep in 'unused' 
               #display; also used as a lower threshold in peptide annotation
plotRes= 300 # dpi
#################################################


rownames(expDes2)= samp_names
colnames(expDes2)=conds
expDes2

samp_vec= rep(rownames(expDes2),each=dim(expDes2)[2])
lab_vec= as.vector(t(expDes2))
expDes2_vec= apply( cbind.data.frame(samp_vec,lab_vec),1,function(x){paste(x[1],x[2],sep='_')})
expDes2_vec_forSites= apply( cbind.data.frame(samp_vec,lab_vec),1,
                             function(x){paste(x[2],x[1],sep='.')})
labs <- apply(cbind.data.frame(rep(samp_names,each=length(conds)),rep(conds,times=length(samp_names))),1,
              function(x){paste(x,collapse='_')})

  if(grepl("\\|.*\\|",protein_oI)) {
    protein_oI_short= str_split(str_split(protein_oI,"\\|")[[1]][3],"_")[[1]][1]  #e.g. get "NFAC4" our of "sp|Q14934|NFAC4_HUMAN"
  } else {
    protein_oI_short= protein_oI
  }

##################################################


#function to create 'sequence lable' for plot
#takes isoform sequences as input
#returns three strings: sequence before relevant p-site, relevant p-site, string after relevant p-site
create_lable= function(isoforms, posInPep.evid,isofGr,plotNo_horiz) 
{
  ##create modified ppep with max. localization probability at each position
   #note that the input isoforms may have brakets indicating localization probabilities at
    #different postions
  
   # 1, find at which AA-positions in the peptides the localization probabilities occur
  sites_wLoc= c()  #vector of positions of sites with localization probability
  for (iso in isoforms ) {
    curr_seq= iso
      while(grepl("\\(", curr_seq)) {
      x= str_locate(curr_seq,"\\([01]\\.?\\d*\\)")  #find first occurrence of loc. prob. in string
      sites_wLoc= c(sites_wLoc,x[[1]][1]-1)
      curr_seq= str_replace(curr_seq,"\\([01]\\.?\\d*\\)","") #remove the first occurence of loc. prob. in string
      }
  }
  sites_wLoc= sort(unique(sites_wLoc))
  sites_wLoc
  
   # 2, Put the corresponding probabilities into array [rows= isoform; columns= site in pep]
  locProb_mtx= matrix(nrow= length(isoforms),ncol= length(sites_wLoc))
  for (isoCount in 1:length(isoforms) ) {
    curr_seq2= isoforms[isoCount]
    for(siteCount in 1:length(sites_wLoc)) {
      site= sites_wLoc[siteCount]
      curr_subSeq2= substr(curr_seq2,site+1,nchar(curr_seq2))  #sub-sequence w/o AA before site prob of interest
        if (grepl("^\\(",curr_subSeq2)) { #string has the loc prob
          st_end = str_locate(curr_subSeq2,"^\\([01]\\.?\\d*\\)")
          locProb_mtx[isoCount,siteCount]= as.numeric( substr(curr_subSeq2,2,st_end[2]-1) )
          curr_seq2= str_replace(curr_seq2,"\\([01]\\.?\\d*\\)","") #remove the first occurence of loc. prob. in string
        } else { # string does not have probability at start
          locProb_mtx[isoCount,siteCount]=  0 }
    }
  }
  locProb_mtx= round(locProb_mtx,2)
  locProb_mtx
  max_locProb_vec= apply(locProb_mtx,2,max)
  
  # 3, Create the peptide sequence with max. loc. probs
  baseSeq= str_replace_all(isoforms[1],'\\([01]?\\.?\\d+\\)','')
  sitesAndEnd= c(sites_wLoc,nchar(baseSeq))  #positions of sites and last letter
  modPep_wMax= substr(baseSeq,1,sitesAndEnd[1])
  for (s in 1:(length(sitesAndEnd)-1)){
    modPep_wMax= paste(modPep_wMax,"(",max_locProb_vec[s],")",
                       substr(baseSeq,sitesAndEnd[s]+1,sitesAndEnd[s+1]),sep='')
  }
  modPep_wMax
  
  #find the position in modified peptide (i.e. entry of all_phosProt_Frac_z):
  #split the string, then count letters only until you reach position of phosphate in peptide (posInPep.STYsites)
  #at the same time count all characters (not just letters)
  modPep_chars= (str_split(modPep_wMax,''))[[1]]   #e.g. split FGLGS(1)PLPS(1)PR
  letter_count=0
  char_count= 0
  while (letter_count< posInPep.evid) { #compare to position of peptide in evidence.txt
    char_count= char_count+1
    #print (modPep_chars[char_count])
    if (grepl("[A-Z]",modPep_chars[char_count])) {  #strictly should only be AA-characters
      letter_count= letter_count+1
    }
  }
  
  
  pre= site_oI-posInPep.evid+1
  xxx=substr(modPep_wMax,1,char_count-1)
  yyy=substr(modPep_wMax,char_count,char_count)
  zzz=substr(modPep_wMax,char_count+1,nchar(modPep_wMax))
  AAinPep= length(str_match_all(modPep_wMax,"[A-Z]")[[1]])  #modPep_wMax e.g. FGLGS(1)PLPS(1)PR
  suff= pre+AAinPep-1
  
  # 4, Find the non-phosphate modifications in the peptide
  mods= (str_split(isofGr,'_'))[[1]][2]   # isofGr e.g. MMYSAACR_Acetyl (Protein N-term),Oxidation (M),Phospho (STY)"
    if (str_detect(mods,',\\d+ Phospho \\(.*\\),?')) {   #e.g. 'Acetyl (Protein N-term), 2 Phospho (STY)' 
      str1= str_split(mods,' Phospho \\(.*\\)')[[1]][1]
      no_phos= as.numeric(str_extract(str1,'\\d+$'))
      mods1= str_replace(mods,'\\d+ Phospho \\(',paste('+',no_phos-1,' Phospho \\(',sep=''))
    } else if (str_detect(mods,'\\d+ Phospho \\(.*\\),?')) { #e.g. '2 Phospho (STY)'
      no_phos= as.numeric(str_extract(mods,'^\\d+'))
      mods1= str_replace(mods,'\\d+ Phospho \\(',paste('+',no_phos-1,' Phospho \\(',sep=''))
    } else {
      mods1= str_replace(mods,',?Phospho \\(.*\\),?','')
    }
    
  # lab= list(bquote(.(protein_oI)*" site:"*.(site_oI)),
  #           bquote(.(xxx)*bold(.(yyy))*.(zzz)),
  #           bquote(.(mods1)) )
  sites_inProt= site_oI-posInPep.evid+sites_wLoc  #calculate sites in protein
  
  total_char_woMod= nchar(pre)+nchar(xxx)+nchar(yyy)+nchar(zzz)+nchar(suff)
  total_char= nchar(pre)+nchar(xxx)+nchar(yyy)+nchar(zzz)+nchar(suff)+nchar(mods1)
  maxChar_perLine= NA
  if(total_char_woMod/plotNo_horiz < 45) {  #don`t split peptide over lines
    maxChar_perLine= max(nchar(paste(protein_oI," site:",site_oI,sep='')),
                         nchar(paste(pre,xxx,yyy,zzz,suff,sep='')),
                         nchar(mods1) )
    lab_out= list(bquote(.(protein_oI)*" site:"*.(site_oI)),
                         bquote( ""^.(pre)*.(xxx)*bold(.(yyy))*.(zzz)*""^.(suff) ),  #I didn`t get subscript working for pre and suff..
                         bquote(.(mods1)),
                  sites_inProt,
                  max_locProb_vec,
                  maxChar_perLine)}  else {
    #split peptide over lines
    if(nchar(xxx) > total_char/2) {
      xxx1= substr(xxx,1,total_char/2)
      xxx2= substr(xxx,(total_char/2)+1,nchar(xxx))
       #avoid splitting at parenthesis
      if(substr(xxx1,total_char/2,total_char/2)=="(") {
        xxx1= substr(xxx,1,total_char/2-1)
        xxx2= substr(xxx,(total_char/2),nchar(xxx))
      } else if (substr(xxx2,1,1)==")") {
        xxx1= substr(xxx,1,total_char/2+1)
        xxx2= substr(xxx,(total_char/2)+2,nchar(xxx))
      }
      maxChar_perLine= max(nchar(paste(protein_oI," site:",site_oI,sep='')),
                           nchar(paste(pre,xxx1,sep='')),
                           nchar(paste(xxx2,yyy,zzz,suff,mods1,sep='')) )
      lab_out= list(bquote(.(protein_oI)*" site:"*.(site_oI)),
                    bquote( ""^.(pre)*.(xxx1) ),  #I didn`t get subscript working for pre and suff..
                    bquote(.(xxx2)*bold(.(yyy))*.(zzz)*""^.(suff)*" "*.(mods1)),
                    sites_inProt,
                    max_locProb_vec,
                    maxChar_perLine)
    } else {
    splitPnt_frEnd= total_char/2 - nchar(mods1)
    splitPnt_zzz= nchar(zzz)-nchar(suff)-splitPnt_frEnd
    zzz1= substr(zzz,1,splitPnt_zzz)
    zzz2= substr(zzz,splitPnt_zzz+1,nchar(zzz))
    #avoid splitting at parenthesis
    if(substr(zzz1,splitPnt_zzz,splitPnt_zzz)=="(" & splitPnt_zzz>1) {
      zzz1= substr(zzz,1,splitPnt_zzz-1)
      zzz2= substr(zzz,splitPnt_zzz,nchar(zzz))
    } else if (substr(zzz2,1,1)==")") {
      zzz1= substr(zzz,1,splitPnt_zzz+1)
      zzz2= substr(zzz,splitPnt_zzz+2,nchar(zzz))
    }
    maxChar_perLine= max(nchar(paste(protein_oI," site:",site_oI,sep='')),
                         nchar(paste(pre,xxx,yyy,zzz1,sep='')),
                         nchar(paste(zzz2,suff,mods1,sep='')) )
    lab_out= list(bquote(.(protein_oI)*" site:"*.(site_oI)),
                  bquote( ""^.(pre)*.(xxx)*bold(.(yyy))*.(zzz1) ),  #I didn`t get subscript working for pre and suff..
                  bquote( .(zzz2)*""^.(suff)*" "*.(mods1)),
                  sites_inProt,
                  max_locProb_vec,
                  maxChar_perLine)
    }
  }
  return(lab_out)
}
##########################################################################
##########################################################################
siteShift= function(baseSeq.isofGr,baseSeq.STYsites) {
  #find how base sequence of isofGr compares to phospho(STY)sites base sequence
  pos_isofGr_in_STYsites= 'NA'
  first4.isofGr= paste(str_split(baseSeq.isofGr,'')[[1]][1:min(4,nchar(baseSeq.isofGr))],collapse='')
  first4.STYsites= paste(str_split(baseSeq.STYsites,'')[[1]][1:min(4,nchar(baseSeq.STYsites))],collapse='')
  #case 1: sequene in isofGr is subsequence of that in STYsites
    if (gregexpr(pattern= baseSeq.isofGr,baseSeq.STYsites) [[1]][1]> 0) {
    pos_isofGr_in_STYsites= -1*(gregexpr(pattern= baseSeq.isofGr,baseSeq.STYsites) [[1]][1]-1)
    #subtracting 1 as no shift will be made if same start site (==1)
    #multiplying with -1 as I will need to shift left from pos in STY-sites
    #case 2: sequence of STYsites is subsequence of that in isofGr
  } else if (gregexpr(pattern= baseSeq.STYsites, baseSeq.isofGr) [[1]][1]> 0){
    pos_isofGr_in_STYsites= gregexpr(pattern= baseSeq.STYsites, baseSeq.isofGr) [[1]][1]-1
    #case 3: sequence in isofGr starts later and finishes later than in STYsites:
    #look only for the first four characters of isofGr
  } else if (gregexpr(pattern= first4.isofGr,baseSeq.STYsites) [[1]][1] != -1) {#string found
      if (length(gregexpr(pattern= first4.isofGr,baseSeq.STYsites) [[1]])> 1)
    { #multiple matches in string
      stop("Cannot match",baseSeq.isofGr,"and",baseSeq.STYsites,"unambiguously!")
    } else {
      pos_isofGr_in_STYsites= -1*(gregexpr(pattern= first4.isofGr,baseSeq.STYsites) [[1]][1]-1)
    }
    #case 4: sequence in STYsites starts later and finishes later than in isofGr:
    #look only for the first four characters of STYsites
  } else if (gregexpr(pattern= first4.STYsites,baseSeq.isofGr) [[1]][1] != -1) {#string found
      if (length(gregexpr(pattern= first4.STYsites,baseSeq.isofGr) [[1]])> 1)
    { #multiple matches in string
      stop("Cannot match",baseSeq.isofGr,"and",baseSeq.STYsites,"unambiguously!")
    } else {
      pos_isofGr_in_STYsites= gregexpr(pattern= first4.STYsites,baseSeq.isofGr) [[1]][1]-1
    }
  } else {
    stop("Cannot match",baseSeq.isofGr,"and",baseSeq.STYsites,"unambiguously!")
  }
  return(pos_isofGr_in_STYsites)
}
 
####################################################
####################################################
main_func= function(evid,evidIDs,suffix,posInPep.STYsites,baseSeq.STYsites,   #seqWprob.STYsites,
                    minLoc) {
  print(paste('MQ evidIDs:',paste(evidIDs,collapse=' ')))
  
  evid_rowNos= match(evidIDs,evid$id)  #row numbers in evidence.txt
  evid_SoI= evid[evid_rowNos,] #evidence for site of interest
  dim(evid_SoI)
  evid_SoI$Phospho..STY..Probabilities
  evid_SoI$Experiment
  evid_SoI$Fraction
  evid_SoI$isofGr= paste(evid_SoI$Sequence,evid_SoI$Modifications,sep='_') #isofrom groups
  
  uniq_isofGr= unique(evid_SoI$isofGr)
  isofGr_count= 0
  
  #globMax= max( apply(evid_SoI[,match(c('Intensity.L','Intensity.M','Intensity.H'),colnames(evid_SoI))],
  #     1,max) )
  
  
  int_vec_sum= rep(0, length(expDes2_vec)) #sum of all intensities for current site
  suppressed_unused_plots= 0 #number of plots for 'unused' group that are not produced
    #because no feature or MSMS meets minimum localization probability
  for (uifGr in uniq_isofGr){ #"unique isoform groups"
    #get intensities in vector that lists samples in order of 'samp_names' and within each sample in order of conds;
    #i.e. the overall order is as in expDes2_vec
    isofGr_count= isofGr_count+1
    
    baseSeq.isofGr= str_split(uifGr,"_")[[1]][1]
    
    #first get sample name, ... and L, M, H intensity for current isoform group
    curr_DF= evid_SoI[evid_SoI$isofGr== uifGr, 
                      match(c('Type','Experiment','Fraction','Charge','Phospho..STY..Probabilities',
                              'Intensity.L','Intensity.M','Intensity.H','isofGr'),colnames(evid_SoI)) ]
    curr_DF
    
    #get localization probability for site of interest
    phosProb_PC_vec= rep(NA, dim(curr_DF)[1])
    for(lineNo in 1:dim(curr_DF)[1]) {
      if (curr_DF$Phospho..STY..Probabilities[lineNo]== '' && grep('MATCH',curr_DF$Type))   { 
        #no site-probability, because match-between-runs feature
        curr_ProbSeq.evid= NA
      }  else if (curr_DF$Phospho..STY..Probabilities[lineNo] != '') {
        curr_ProbSeq.evid= curr_DF$Phospho..STY..Probabilities[lineNo]
      } else {
        stop("No site-probability, but apparently not a matched feature!")
      }
      
      if(!is.na(curr_ProbSeq.evid)) {
        if (is.na(posInPep.STYsites)) {  #calculate relevant position in curr_ProbSeq.evid from
                                        # site in protein
          posInPep.evid= siteInProt-startPepInProt+1
        } else {
        #find how base sequence of isofGr compares to phospho(STY)sites base sequence
        shift= siteShift(baseSeq.isofGr,baseSeq.STYsites)  
        posInPep.evid= posInPep.STYsites+shift #phospho position in peptide from evidence.txt
        #calculated from site given in phospho(STY)sites.txt and shift between sequences  
        }
        
        curr_ProbSeq.evid_split= str_split(curr_ProbSeq.evid,'')[[1]]  #e.g. split of "IT(0.65)S(0.262)IS(0.044)PT(0.047)PEPPAALEDNPDAWGDGS(0.997)PR"
        char_count=0
        phosProb=NA #if no probability at site of interest -> will remain NA
        for (i in 1:length(curr_ProbSeq.evid_split)){
          if(grepl("[A-Z]",curr_ProbSeq.evid_split[i])) {   #count characters; strictly these should only be AA-letters
            char_count= char_count+1
            #print(paste("char-count:",char_count))
          } else if (grepl("\\(",curr_ProbSeq.evid_split[i]) & char_count== posInPep.evid) {#you arrived at position of mod
            #print(paste("char-count:",char_count))
            siteTillEnd= paste(curr_ProbSeq.evid_split[(i+1):length(curr_ProbSeq.evid_split)],collapse='')
            phosProb= as.numeric(str_extract(siteTillEnd,"^[01]\\.?[0-9]*"))
            break
          } else if (grepl("\\(",curr_ProbSeq.evid_split[i])==FALSE & char_count== posInPep.evid) {
            print(paste("Character after",posInPep.evid,'th letter is',curr_ProbSeq.evid_split[i]))
          }
        print(paste("i:",i,", phosProb:",phosProb))
        }
      } else {
        phosProb= NA }
      phosProb
      phosProb_PC_vec[lineNo]= phosProb*100
    }
    curr_DF$phosProb_PC= phosProb_PC_vec
    
    #remove sequences with too low localization probability
      #only from unused sequences (used ones may have no loc. prob., because they are from
      #match-between-runs)
    if(suffix== 'unused') {
      curr_DF= curr_DF[!is.na(curr_DF$phosProb_PC) & curr_DF$phosProb_PC >= minLoc, ]
        #dont`t allow MbR feqtures, because may be matching to ppep with low loc. prob.
    }
    
    if(sum(!is.na(curr_DF$phosProb_PC) & curr_DF$phosProb_PC >= minLoc)==0){
      print(paste("Isof.Gr.",uifGr,": No localization probability >",minLoc,"!"))
    }
    
    #split DF into MS/MS ('MSMS') and features (all others)
    MSMS_DF= curr_DF[curr_DF$Type == 'MSMS',]
    curr_DF_feat= curr_DF[curr_DF$Type != 'MSMS',]
    
    if (nrow(curr_DF_feat)>0) {    #there are features, not only MS/MS-spectra
      curr_DF_melt_orig= melt(curr_DF_feat,id.vars=c('Experiment','Fraction','Charge','phosProb_PC',
                                        'Phospho..STY..Probabilities'),
                              measure.vars=c('Intensity.L','Intensity.M','Intensity.H'))
      #I do not add 'Phospho..STY..Probabilities' to curr_DF_melt_orig
      colnames(curr_DF_melt_orig)= c('Experiment','Fraction','Charge','phosProb_PC','seqWprob',
                                     'lable','Intensity')
      curr_DF_melt_orig$lable= sapply(curr_DF_melt_orig$lable, function(x){str_replace(x,'Intensity.','')})
      curr_DF_melt_orig$sampLab= apply(curr_DF_melt_orig,1,function(x){paste(x[1],x[6],sep='_')})
      
      curr_DF_melt_a= aggregate(Intensity~Experiment+Fraction+Charge+lable+sampLab,
                                curr_DF_melt_orig,FUN= sum) #in case there are several features for one entity
      curr_DF_melt_b= aggregate(phosProb_PC~Experiment+Fraction+Charge+lable+sampLab,
                                curr_DF_melt_orig,FUN= min,na.action= na.pass)
        #I keep entries with phosProb_PC == NA; these may be from match-between-runs
      curr_DF_melt_c= aggregate(seqWprob~Experiment+Fraction+Charge+lable+sampLab,
                                curr_DF_melt_orig,FUN= min,na.action= na.pass)
      curr_DF_melt_ab= merge.data.frame(curr_DF_melt_a,curr_DF_melt_b,
            by=c('Experiment','Fraction','Charge',
                              'lable','sampLab'))
      curr_DF_melt= merge.data.frame(curr_DF_melt_ab,curr_DF_melt_c,
                                        by=c('Experiment','Fraction','Charge',
                                             'lable','sampLab'))
      
      #Create label for plot below
      isoforms= unique(curr_DF_feat$Phospho..STY..Probabilities[curr_DF_feat$Phospho..STY..Probabilities != ''])
      #excluding empty entries for Phospho..STY..Probabilities, which can arise from match-between-runs features
      #The isofGr column of all entries of curr_DF_feat are expected to be the same
      if(length(unique(curr_DF_feat$isofGr)) > 1) {
        stop('More than one isoform in group!')
      }
      isofGr= curr_DF_feat$isofGr[1]
      
      
      plotNo_horiz= length(unique(curr_DF_melt$Charge))
      plotNo_vert= length(unique(curr_DF_melt$Fraction))
      lab= create_lable(isoforms, posInPep.evid,isofGr,plotNo_horiz)  #plotNo_horiz to decide if to split peptide over rows
      #output: list: prot+site, mod_seq
         #additional mods; vec of pos in pep; vec of probs.
      
      chars= lab[[6]][1] #characters in longest line of main lable
      print(paste("Isof.Gr:",isofGr,"suffix:",suffix,"chars:",chars))
      lab_main= list(lab[[1]],lab[[2]],lab[[3]]) #lable of plot-array

      #Look at each intensity fraction- and charge-wise
      
      #char_size= min(plotNo_horiz/chars*45, 1.2) #determined empirically
      char_size= 1.2 #consistent cex of 1.3, unless...
      if(chars> (40*plotNo_horiz)) {  #if too many characters; there more plots next
                                        #to each other, the more allowed
      char_size= 1.2* 40/(chars/plotNo_horiz)}
      
      #lab_main= list(lab[[1]],lab[[2]],paste(chars,char_size)) #lable of plot-array
      
      png(filename= paste(paste(today,protein_oI_short,site_oI,"isofGr",
          isofGr_count-suppressed_unused_plots,suffix,sep="_"),".png",sep=''),width=length(unique(curr_DF_melt$Charge))*5,
          height=length(unique(curr_DF_melt$Fraction))*5,units="in",res= plotRes)  #plot dimensions depend on number of sub-plots
      par(mfrow=c(plotNo_vert,plotNo_horiz),
          pin= c(4,3.5),oma = c(0, 0, 3, 0),cex=1)  #cex=1 is important to avoid text rescaling
      for (frac in unique(sort(curr_DF_melt$Fraction))){
        for (z in unique(curr_DF_melt$Charge)) {
          curr_DF_Frac_z= curr_DF_melt[curr_DF_melt$Fraction== frac & curr_DF_melt$Charge== z,]
          curr_DF_Frac_z
          
          ##Create label for plot specific to fraction and charge
          isoforms_spec= unique(curr_DF_Frac_z$seqWprob[curr_DF_Frac_z$seqWprob != ''])
          if(length(isoforms_spec)==0) { #not detected in this fraction and charge
            lab_spec= 'ND' 
          } else {
            lab_spec= create_lable(isoforms_spec, posInPep.evid,isofGr,plotNo_horiz)
            sites_prob_spec1= rbind.data.frame(lab_spec[[4]],lab_spec[[5]])
            sites_prob_spec= as.matrix(sites_prob_spec1[,sites_prob_spec1[2,]>minLoc/100],
                nrow=2)  #needs 'as.matrix' otherwise converted to row-vector if only one site
            sites_prob_conc_vec= apply(sites_prob_spec,2,function(x){
                  paste(x[1],': ',x[2],sep='')})
            ##I want to print max. 3 probabilities per row
             ##fill up to multiple of 3
            #toAdd= 3*ceiling(length(sites_prob_conc_vec)/3) - length(sites_prob_conc_vec)
            #sites_prob_conc_vec_filled= c(sites_prob_conc_vec,rep('',toAdd))
            #sites_prob_conc_mtx= matrix(sites_prob_conc_vec_filled,ncol=3,
             #   byrow=TRUE) #not more than 3 per row 
            noProbs_per_row= 3  #number of probabilities to list per row in plot header
            rowsInProbsHeader= ceiling(length(sites_prob_conc_vec)/noProbs_per_row)
            lab_spec= paste(sites_prob_conc_vec[1:min(noProbs_per_row,length(sites_prob_conc_vec))],collapse=', ')
            if(rowsInProbsHeader>1){
              for(rowNo in 2:rowsInProbsHeader){
                lab_spec= paste(lab_spec,
                paste(sites_prob_conc_vec[(((rowNo-1)*noProbs_per_row)+1):
                          min((rowNo*noProbs_per_row),length(sites_prob_conc_vec))],collapse=', '),
                sep= '\n')
              }
            }
            #lab_spec= paste(apply( sites_prob_conc_mtx,1,function(x){paste(x,collapse=' ')}),
             #     collapse='\n')
          #chars_spec= nchar(isoforms_spec[1]) #isoforms at least have approx. same length  
          }
          
          
          int_vec= rep(NA, length(expDes2_vec))
          if(length(curr_DF_Frac_z$phosProb_PC)> 0) {
            if(all(is.na(curr_DF_Frac_z$phosProb_PC))){
              min_phosProb_PC= NA #in case of MBR only feature(s)
            } else {
            min_phosProb_PC= min(curr_DF_Frac_z$phosProb_PC, na.rm= TRUE)}
          } else {
            min_phosProb_PC= NA
          }
          
          for(j in 1:length(expDes2_vec)) {  #for each sample
            if(any(curr_DF_Frac_z$sampLab==expDes2_vec[j])) {  #if there is intensity
              idx= which(curr_DF_Frac_z$sampLab==expDes2_vec[j])  #get its index
              if(length(idx)>1){
                stop('More than one intensity for entity!')
              }
              int_vec[j]= curr_DF_Frac_z$Intensity[idx]
            }
            else {int_vec[j]=0}  #otherwise write 0 for this sample
          }
          print(paste('int_vec:',paste(int_vec,collapse=' ')))
          int_vec_sum= colSums(rbind.data.frame(int_vec,int_vec_sum))
          names(int_vec_sum)= c()
          print(paste('sum: ',paste(int_vec_sum,collapse=' ')))
          
          if(all(is.na(int_vec))) {
            int_vec= rep(0, length(int_vec))
          }
          
          if (same_scale== 'n') {
            y_max= 1.2*max(c(curr_DF_melt$Intensity[curr_DF_melt$Charge== z],10), na.rm= TRUE) #I want all fractions to be
          } else {
            if (length(site_rowNo)> 0){
              y_max=  max(site_ints) } else {
              y_max= 1.2*max(c(curr_DF_melt$Intensity,10), na.rm= TRUE)
            }
          }
          
          print(x<-barplot(int_vec,ylim= c(0,y_max) ))
          
          mtext(do.call(expression,lab_main),side=3,line=c(2.5,0.7,-0.8), 
                outer = TRUE, cex = char_size, padj=1 )
          #same text size for all:
          text_size= 1.55
          
          # if (plotNo_vert ==1) { #the more vertical subplots, the bigger the text
          #   text_size= 1.55 } 
          # else if (plotNo_vert==2) {
          #   text_size= 2.1 }
          # else if (plotNo_vert==3) {
          #   text_size= 2.25  } 
          # else if (plotNo_vert>3) {
          #   text_size= 2.55 }
          ypos= max(0,(max(nchar(labs))/2-1)*(y_max/100))   #empirically determined y-position of column labels
          text(cex=1, x=x, y=ypos, labs, xpd=TRUE, srt=90)
          #text(x=median(x),y=0.97*y_max,paste("   min. loc. prob.",min_phosProb_PC,"%",sep=''),cex=text_size)
          text(x=median(x),y=0.965*y_max,paste('Fraction:',frac,'     charge:',z),cex=text_size)
          text(x=median(x),y=0.86*y_max,lab_spec,cex=text_size)
          
          #print(paste("fraction:",frac,"z:",z,"min. loc. prob.:",min_phosProb_PC))
          }
      }
      dev.off()
    }   else if (nrow(MSMS_DF)>0) {  #only MSMS, no features
      all_phosProb_MSMS= paste(unique(MSMS_DF$Phospho..STY..Probabilities),sep=",") #list of all peptides with unique localization
      #probabilities representing current modified sequence in fraction at given z (I expect that there usually is just one)
      
      png(filename= paste(paste(today,protein_oI_short,site_oI,"isofGr",
          isofGr_count-suppressed_unused_plots,"MSMS",suffix,sep="_"),".png",sep=''),width=5,
          height=5,units="in",res= plotRes)  #plot dimensions depend on number of sub-plots
      par(mfrow=c(1,1),pin= c(4,3.5),oma = c(0, 0, 3, 0),cex=1)  #important to avoid text rescaling)
      print(x<-barplot(rep(0,dim(expDes2)[1]*dim(expDes2)[2]),ylim= c(0,10) ))
      
      #label plot with peptide
      isoforms= unique(MSMS_DF$Phospho..STY..Probabilities)
      #The isofGr column of all entries of curr_DF_feat are expected to be the same
      if(unique(MSMS_DF$isofGr) > 1) {
        stop('More than one isoform in group!')
      }
      isofGr= MSMS_DF$isofGr[1]
      lab= create_lable(isoforms, posInPep.evid,isofGr,1)
      
      mtext(do.call(expression,lab),side=3,line=c(2.5,0.7,-0.8),   
            outer = TRUE, cex = char_size, padj=1 )
      
      #Same text size for all
      text_size= 1.55
      
      # if (plotNo_vert ==1 ) { #the more vertical subplots, the bigger the text
      #   text_size= 1.1 } 
      # else if (plotNo_vert==2) {
      #   text_size= 1.4 }
      # else if (plotNo_vert==3) {
      #   text_size= 1.5  } 
      # else if (plotNo_vert>3) {
      #   text_size= 1.7 }
      ypos= max(0,max(nchar(labs))/2-1)   #empirically determined y-position of column labels
      text(x=median(x),y=5,"No intensities recorded",cex=text_size)
      text(cex=1, x=x, y=ypos, labs, xpd=TRUE, srt=90)
      
      dev.off()
    } else {
      suppressed_unused_plots=+ 1
    }
    
  }
  #Plot summed evidence intensities (should equal site-intensities)
  png(filename= paste(paste(today,protein_oI_short,site_oI,"site",suffix,"_calc",sep="_"),".png",sep=''),
      width=5, height=5, units="in",res= plotRes)  #plot dimensions depend on number of sub-plots
    if (suffix == 'unused') {
      if (same_scale== 'y') {
        print(x<-barplot(int_vec_sum, col='blue',ylim=c(0,max(site_ints)))) } else {
        print(x<-barplot(int_vec_sum, col='blue'))
        }
    } else {
      if (same_scale== 'y') {
        print(x<-barplot(int_vec_sum, col='green',ylim=c(0,max(site_ints)))) } else {
          print(x<-barplot(int_vec_sum, col='green'))
        }
    }
  text(cex= 1, x=x, y=0, labs, xpd=TRUE, srt=90)
  mtext( bquote(atop(.(protein_oI)*" site:"*.(site_oI))),cex = 1.2)
  dev.off()
}
####################################################
####################################################

#I/O
setwd(file_location)
sites= read.csv('Phospho (STY)Sites.txt',sep='\t',header=TRUE, stringsAsFactors = FALSE)
dim(sites)

evid= read.csv('evidence.txt',sep='\t',header = TRUE, stringsAsFactors = FALSE)
dim(evid)

# Plot intensities from "phospho(STY) sites.txt"
IntVec_forSites= paste(rep("Intensity",length(expDes2_vec_forSites)),expDes2_vec_forSites,sep='.')
intIdx_sites= match(IntVec_forSites,colnames(sites))
site_rowNo= which(sites$Protein==protein_oI & 
  (grepl(paste("^",site_oI,"$",sep=''),sites$Positions.within.proteins) |
  grepl(paste("^",site_oI,";",sep=''),sites$Positions.within.proteins)) )   #row number in sites-file
 #'Position.within.proteins' may have several entries, if isoforms -> look at first before ';'

if (length(site_rowNo)> 0) {
  site_ints= as.numeric(sites[site_rowNo,intIdx_sites])
  
  png(filename= paste(paste(today,protein_oI_short,site_oI,"site",sep="_"),".png",sep=''),
    width=5, height=5, units="in",res= plotRes)  #plot dimensions depend on number of sub-plots
  print(x<-barplot(site_ints, col='red'))
  text(cex= 1, x=x, y=0, labs, xpd=TRUE, srt=90)
  mtext( bquote(atop(.(protein_oI)*" site:"*.(site_oI))),cex = 1.2)
  dev.off()
}

#Look up corresponding evidences
if (length(site_rowNo)==0) {
  evidIDs= c()
  print(paste("Site",site_oI,"on protein",protein_oI,"not found!"))
} else {
  evidID_str= sites$Evidence.IDs[site_rowNo]  #semi-colon separated entries
  evidIDs= as.numeric(str_split(evidID_str,';')[[1]])
  
  posInPep.STYsites= sites$Position.in.peptide[site_rowNo]  #position in peptide in phospho(STY)sites.txt
  seqWprob.STYsites= sites$Phospho..STY..Probabilities[site_rowNo]  #e.g. TSQLT(0.002)DHS(0.998)K
  baseSeq.STYsites= str_replace_all(seqWprob.STYsites,'\\(\\d+\\.?\\d*\\)','') #e.g. TSQLTDHSK
  
  suffix= ''
  startPepInProt= NA
  siteInProt= site_oI
  main_func(evid, evidIDs,suffix,posInPep.STYsites,baseSeq.STYsites,minLoc)   #seqWprob.STYsites,
  #main_func also considers evidences other than with 'baseSeq', but 'baseSeq' and 
  #'posInPep' are needed to calculate position in other ppeps
}

####################
# Plot label-ratios from "phospho(STY) sites.txt"
if (length(site_rowNo)> 0) {
  expDes2
  
  ratio_mtx= data.frame(matrix(NA,nrow=nrow(expDes2),ncol=ncol(expDes2)-1)) #rows are samples; 
     #columns are comparisons 2-vs-1 and 3-vs-2
  rownames(ratio_mtx)= samp_names
  colnames(ratio_mtx)= c("2vs1","3vs2")
  for (i in 1:dim(expDes2)[1]) {
    comp2to1= paste("Ratio",expDes2[i,2],expDes2[i,1],"normalized",rownames(expDes2)[i],
                    sep='.')   #e.g. "Ratio M/L normalized R1"
    if ( is.na(match(comp2to1, colnames(sites)))) {  #ratio doesn`t exist -> look for inverse
      comp1to2= paste("Ratio",expDes2[i,1],expDes2[i,2],"normalized",rownames(expDes2)[i],
                      sep='.')
      if (is.na(match(comp1to2, colnames(sites)))) {
        print (paste("Error: Neither", comp2to1, "nor", comp1to2, "found"))
      } else {
        ratio_mtx[i, 1]= 1/sites[site_rowNo, match(comp1to2,colnames(sites))]
      }
    } else {
      ratio_mtx[i, 1]= sites[site_rowNo, match(comp2to1, colnames(sites))]
    } 
    
    comp3to2= paste("Ratio",expDes2[i,3],expDes2[i,2],"normalized",rownames(expDes2)[i],
                    sep='.')   #e.g. "Ratio M/L normalized R1"
    if ( is.na(match(comp3to2, colnames(sites)))) {  #ratio doesn`t exist -> look for inverse
      comp2to3= paste("Ratio",expDes2[i,2],expDes2[i,3],"normalized",rownames(expDes2)[i],
                      sep='.')
      if (is.na(match(comp2to3, colnames(sites)))) {
        print (paste("Error: Neither", comp3to2, "nor", comp2to3, "found"))
      } else {
        ratio_mtx[i, 2]= 1/sites[site_rowNo, match(comp2to3,colnames(sites))]
      }
    } else {
      ratio_mtx[i, 2]= sites[site_rowNo, match(comp3to2, colnames(sites))]
    } 
  }
  ratio_mtx
  ratio_mtx$samp= as.factor(rownames(ratio_mtx))
  ratio_mtx_melt= melt(ratio_mtx, id.vars="samp")   #measure.vars = c("2vs1","3vs2"))
  limit= max(max(ratio_mtx_melt$value,na.rm = TRUE), 1/min(ratio_mtx_melt$value,na.rm=TRUE),2)
  
  labs_ratio= rep(NA,dim(expDes2)[2]-1)  #create labels for ratio-plot
  for(s in 1:dim(expDes2)[2]-1){
    labs_ratio[s]= paste(colnames(expDes2)[s+1],'/',colnames(expDes2)[s])
  }
  png(filename= paste(paste(today,protein_oI_short,site_oI,"site_MQratios",sep="_"),".png",sep=''),
      width=5, height=5, units="in",res= plotRes)  #plot dimensions depend on number of sub-plots
  print(x<- ggplot(data= ratio_mtx_melt, aes(x= samp,y=value,group=variable))+
          geom_bar(stat="identity",position=position_dodge(),aes(fill=variable),colour="black")+
          scale_x_discrete(name='sample')+
          scale_y_continuous(name='MaxQuant-Norm. ratio', trans="log2",limits= c(1/limit,limit),
                             breaks=c(0.2,0.33,0.4,0.5,0.67,0.83,1,1.2,1.5,2,2.5,3,4) )+
          scale_fill_manual(name="",
                            values=c("#999999", "#E69F00"),
                            labels= labs_ratio)+
          #guides(fill=FALSE)+
          ggtitle(paste(protein_oI,", site:",site_oI,sep=''))
  )
  dev.off()
} else {
  png(filename= paste(paste(today,protein_oI_short,site_oI,"site_MQratios",sep="_"),".png",sep=''),
      width=5, height=5, units="in",res= plotRes)  #plot dimensions depend on number of sub-plots
  print( 
  plot(x=c(0,1),y=c(0,1),type="n",axes= FALSE,xlab='',ylab=''))
  text(0.5,0.5,"No site-entry for this phospho-site") 
  dev.off()
}


###########################################
# evidences not used by MQ: find evidences from protein_oI that cover site_oI, but were not 
  #used for p-Site quantification
 #read .fasta
fastaFile <- readAAStringSet(fasta)
fasta_seqName = names(fastaFile)
fasta_seqVec = paste(fastaFile)

seq_oI= fasta_seqVec[sapply(fasta_seqName, function(x) 
       {str_detect(x, paste('\\|',protein_oI,'\\|',sep=''))})]
if(length(seq_oI)==0){
  print("Protein sequence not found. Did you select correct .fasta file?")}

#find evidences of protein of interest in evidence.txt
search_prot= paste("(^|;)",str_replace_all(protein_oI,"\\|","\\\\|"),"($|;)",sep='') #generate search string to search for protein_oI
proI_eviIDX =which(str_detect(evid$Proteins,search_prot ))
evid$Proteins[proI_eviIDX]

#limit to evidences of phospho-peptides
length(evid$Modifications[proI_eviIDX])
proI_phosEviIDX= proI_eviIDX[ evid$Phospho..STY.[proI_eviIDX] > 0]
length(proI_phosEviIDX)
  

#limit to evidences that have intensity, i.e. not only MSMS
proI_phosFeatIDX= proI_phosEviIDX[evid$Type[proI_phosEviIDX] != "MSMS"]
length(proI_phosFeatIDX)
    
#limit to evidences that are not used for p-site by MQ (i.e. not among evidence IDs)
proI_phosFeatIDs= evid$id[proI_phosFeatIDX]
unused_FeatIDs= proI_phosFeatIDs[which(is.na(match(proI_phosFeatIDs,evidIDs)))]

if(length(unused_FeatIDs)>0) {
    #determine if sequence of evidences includes site-of-interest
    unusedEvid_seqs= evid$Sequence[ match(unused_FeatIDs, evid$id)]
    unusedEvid_pos= str_locate_all(seq_oI,unusedEvid_seqs)
    unused_oI_IDs= c()
    unused_oI_startPos= c()
    for (i in 1:length(unusedEvid_pos)) {
      for (j in 1:(dim(unusedEvid_pos[[i]])[1])) {   #dim1: how often string was found; dim2= 2 (start,end)
        if (site_oI >= unusedEvid_pos[[i]][j,1] && site_oI <= unusedEvid_pos[[i]][j,2]) {
          unused_oI_IDs= c(unused_oI_IDs,unused_FeatIDs[i])
          startPepInProt= unusedEvid_pos[[i]][j,1]
          unused_oI_startPos= c(unused_oI_startPos,startPepInProt)
          break
        }
      }
    }
    unused_siteEvid= evid[match(unused_oI_IDs,evid$id),]
    unused_siteEvid
    
    if(dim(unused_siteEvid)[1] > 0){
      seqWprob= unused_siteEvid$Phospho..STY..Probabilities[1]  #
      baseSeq= unused_siteEvid$Sequence[1]   #
      
      suffix= 'unused'
      startPepInProt= unused_oI_startPos[1] #the first of the unused evidence peptides represents
        #the site (position of phosphate of interest)
      posInPep= site_oI- (startPepInProt-1)
      #I will only plot unused evidences above min. loc. prob.
      main_func(unused_siteEvid,unused_siteEvid$id,suffix,posInPep,baseSeq,minLoc) #seqWprob,
       #main_func also considers evidences other than with 'baseSeq', but 'baseSeq' and 
         #'posInPep' are needed to calculate position in other ppeps
  }
}
```
